# Responses of globally important phytoplankton species to olivine dissolution products and implications for carbon dioxide removal via ocean alkalinity enhancement

**DOI:** 10.1101/2023.04.08.536121

**Authors:** David A. Hutchins, Fei-Xue Fu, Shun-Chung Yang, Seth G. John, Stephen J. Romaniello, M. Grace Andrews, Nathan G. Walworth

## Abstract

Anthropogenic greenhouse gas emissions are leading to global temperature increases, ocean acidification, and significant ecosystem impacts. Given current emissions trajectories, the IPCC reports indicate that rapid abatement of CO_2_ emissions and development of carbon dioxide removal (CDR) strategies are needed to address legacy and difficult to abate emissions sources. These CDR methods must efficiently and safely sequester gigatons of atmospheric CO_2_. Coastal Enhanced Weathering (CEW) via the addition of the common mineral olivine to coastal waters is one promising approach to enhance ocean alkalinity for large-scale CDR. As olivine weathers, it releases several biologically active dissolution products, including alkalinity, trace metals, and the nutrient silicate. Released trace metals can serve as micronutrients but may also be toxic at high concentrations to marine biota including phytoplankton that lie at the base of marine food webs. We grew six species representing several globally important phytoplankton species under elevated concentrations of olivine dissolution products via a synthetic olivine leachate (OL) based on olivine elemental composition. We monitored their physiological and biogeochemical responses, which allowed us to determine physiological impacts and thresholds at elevated olivine leachate concentrations, in addition to individual effects of specific constituents. We found both positive and neutral responses but no evident toxic effects for two silicifying diatoms, a calcifying coccolithophore, and three cyanobacteria. In both single and competitive co-cultures, silicifiers and calcifiers benefited from olivine dissolution products like iron and silicate or enhanced alkalinity, respectively. The non-N_2_-fixing picocyanobacterium could use synthetic olivine-derived iron for growth, while N_2_-fixing cyanobacteria could not. However, other trace metals like nickel and cobalt supported cyanobacterial growth across both groups. Growth benefits to phytoplankton groups *in situ* will depend on species-specific responses and ambient concentrations of other required nutrients. Results suggest olivine dissolution products appear unlikely to cause negative physiological effects for any of the phytoplankton examined, even at high concentrations, and may support growth of particular taxa under some conditions. Future studies can shed light on long-term eco-evolutionary responses to olivine exposure and on the potential effects that marine microbes may in turn have on olivine dissolution rates and regional biogeochemistry.

## Introduction

Excess anthropogenic greenhouse gas emissions are driving global changes to Earth systems and leading to simultaneous increases in sea surface temperatures, ocean acidification, and regional shifts in nutrient supplies (IPCC, 2022). To counteract these trends and limit the average global temperature increase to 1.5-2°C, carbon dioxide removal (CDR) methods that can collectively remove and permanently store gigatons of atmospheric CO_2_ (GtCO_2_) must be developed (Rogelj et al., 2018). Coastal Enhanced Weathering (CEW) with olivine (Mg_2-x_Fe_x_SiO_4_) has been proposed as an economically scalable form of ocean alkalinity enhancement (OAE), as it is a globally abundant, naturally occurring ultramafic silicate mineral (Taylor et al., 2016; Caserini et al., 2022). Olivine is considered to be one of the most favorable minerals for CDR as it weathers quickly under Earth surface conditions (Oelkers et al., 2018). Like other silicate minerals, it dissolves in water to release cations (Mg^2+^, Fe^2+^) and generates alkalinity (principally HCO_3_^−^), with up to 4 mol of CO_2_ sequestered per mol of olivine [Eq. 1].

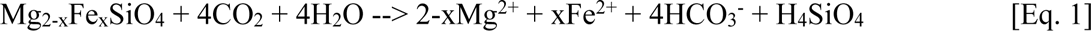

Forsteritic olivine is the magnesium-rich end-member of olivine and can contain various other trace constituents. For example, olivine used in this study contains ∼92% magnesium (Mg^2+^) and ∼8% ferrous iron (Fe^2+^) along with trace amounts (<1%) of other metals such as nickel (Ni), chromium (Cr), and cobalt (Co). As olivine weathers, it releases several biologically important dissolution products into the surrounding seawater: (I) bicarbonate (HCO_3_^−^) and carbonate ion (CO_3_^2−^), hereafter summarized as “alkalinity”; (II) silicic acid (Si(OH)_4_) hereafter termed silicate; (III) and a variety of trace metals including iron (Fe^2+^, or oxidized aqueous species), nickel (Ni^2+^), cobalt (Co^2+^), and chromium (CrVI). These dissolution products have the potential to affect important phytoplankton functional groups like silicifying algae (diatoms), calcifying algae (coccolithophores), and cyanobacteria, which lie at the base of marine food webs and drive the biological carbon pump (Hauck et al., 2016; Moran, 2015). Hence, it is important to understand the specific effects of these constituents on globally important phytoplankton species, particularly at elevated concentrations to simulate large-scale CEW applications.

Significant alkalinity additions from olivine weathering can consume CO_2_ from the surrounding seawater, causing a CO_2_ deficit until air-sea equilibration. This shift in the carbonate system from CO_2_ to HCO_3_^−^/CO_3_^2−^ by transient, non-equilibrated OAE may affect phytoplankton functional groups differently, with some taxa being more sensitive than others. For example, it is predicted that calcifying organisms like coccolithophores may benefit from CEW due to decreases in proton concentrations (H^+^) and increases in the CaCO_3_ saturation state. Additionally, dissolving one mole of olivine leads to a one mole increase in dissolved silicate, which is an essential and often bio-limiting nutrient for silicifying organisms like diatoms, a phytoplankton group estimated to contribute up to 40% of the marine primary production (Bertrand et al., 2012). Hence, diatoms may especially benefit from CEW applications with olivine. Additionally, diatoms are particularly noted for being dominant phytoplankton in the coastal regimes where olivine deployments are likely to take place (Field et al., 1998). While there are both planktonic and benthic species of diatoms, the latter will presumably be exposed to especially sustained and elevated levels of dissolution products when olivine is deployed in natural marine sediments. It is unknown if either group, calcifiers or silicifiers, may consistently outcompete the other following CEW with olivine (Bach et al., 2019).

Trace metals like Fe and Ni are general micronutrients required by all classes of phytoplankton and could potentially support their growth upon fluxes into seawater from olivine weathering. In particular, dinitrogen (N_2_)-fixing cyanobacteria and diatoms both have elevated Fe requirements (Hutchins and Sañudo-Wilhelmy, 2021; Hutchins and Boyd, 2016), and so may stand to benefit from increases in Fe concentrations. Although a required micronutrient at low levels, in high enough concentrations Ni may potentially negatively impact phytoplankton growth, although one recent study showed limited to no toxic effects of very high Ni concentrations (e.g. 50,000 nmol L^−1^) for several phytoplankton taxa (Guo et al., 2022). Cobalt can also serve as a micronutrient for phytoplankton (Sunda and Huntsman, 1995; Hawco et al., 2020) but may also be toxic at high concentrations (Karthikeyan et al., 2019). However, other trace metals found with olivine such as Cr are not nutrient elements and also need to be considered in terms of their possible toxicity to phytoplankton (Flipkens et al., 2021; Frey et al., 1983).

Hence, it is important to understand the taxon-specific effects of these constituents to determine thresholds at which key phytoplankton functional groups may experience positive or negative effects. Furthermore, it is important to expose phytoplankton to elevated concentrations of olivine dissolution products simultaneously to understand what impacts may occur for large CEW applications. Exposures of organisms to concentrated olivine dissolution products also provides an “worst case scenario” benchmark, which can be compared to lower actual environmental exposures resulting from small CEW additions, slower olivine dissolution time scales, and dilution from advection. While olivine weathers relatively quickly compared to other silicate minerals (Hartmann et al., 2013), dissolution of olivine grains is gradual (i.e. years) relative to microbial physiological responses (hours), posing a challenge to test different concentrations of olivine constituents on phytoplankton physiology. To address this, we prepared a synthetic olivine leachate (OL) composed of olivine dissolution products with trace metal concentrations well over those of seawater (7-12,000 times higher), in order to represent a “worst case” scenario for a CEW project. This extreme scenario was estimated based on the maximum expected impact of olivine weathering on the chemistry of the overlying water column. Assuming a 10 cm thick layer of pure olivine sand dissolves with a 100 year half-life into 1 meter of overlying water with a 24 hour residence time, the anticipated steady state change in the alkalinity of the overlying water column is 65 umol/kg (assuming 4 moles of alkalinity per mole olivine (Meysman and Montserrat, 2017) and 100% release to the water column). The concentrations of other components were chosen assuming stoichiometric, congruent dissolution and quantitative release to the water column as well -- a worst case scenario. Furthermore, phytoplankton were exposed to OL within a small, enclosed batch culture. We cultured 6 species representing three globally important phytoplankton functional groups: 2 diatoms (*Nitzschia, Ditylum*), 1 coccolithophore (*Emiliana*), 2 dinitrogen (N_2_) fixing cyanobacteria (*Trichodesmium*, *Crocosphaera*), and 1 non-N_2_ fixing picocyanobacterium (*Synechococcus*). All of these species are planktonic, with the exception of the diatom *Nitzschia* which frequently forms benthic biofilms (Yamamoto et al., 2008). Cultures were grown semi-continuously in natural seawater based modified Aquil media (Sunda et al., 2005) with OL as the only available Fe source (and Si source for diatoms). For all experiments, cultures were sampled for a basic set of core biogeochemical and physiological parameters (Fu et al., 2005, 2008; Tovar-Sanchez et al., 2003; Paasche et al., 1996). This approach allowed us to compare phytoplankton taxon-specific responses, including: **1)** physiological impacts at extremely high OL concentrations, **2)** physiological thresholds and dose responses across a range of increasing concentrations of OL, and **3)** individual effects of specific OL constituents.

## Materials and Methods

### Culture growth conditions and experimental set up

Six species of phytoplankton were used in these experiments, including: the planktonic diatom *Ditylum brightwellii* (centric, planktonic, isolated by T. Rynearson from Narragansett Bay, Rhode Island, USA) and the benthic diatom *Nitzschia punctata* (CCMP 561, isolated from tidal mud near San Diego, North Pacific Ocean), a coccolithophore, *Emiliania huxleyi (*CCMP 371, a North Atlantic isolate), a picoplanktonic cyanobacterium *Synechococcus sp.* (strain XM-24, isolated by J. Zheng from Xiamen estuary, the South China Sea, PRC, belonging to clade CB5, subcluster 5.2), and two marine dinitrogen (N_2_) fixing cyanobacteria, *Trichodesmium erythraeum* (strain IMS 101, from the Gulf Stream, Northwest Atlantic Ocean) and *Crocosphaera watsonii* (WH 0005, from the North Pacific Ocean). Cultures were grown in 500 mL polycarbonate flasks at 28^0^C for the three cyanobacteria, and 20^0^C for the diatoms and *Emiliania huxleyi*. Cool-white fluorescent light was supplied following a 12:12 light:dark cycle at an irradiance level of 150 µEm^−2^s^−1^. Stock cultures were grown in natural offshore seawater collected with trace metal clean methods (John et al., 2022), which was used to make modified Aquil Control Medium (ACM). The positive control ACM contained replete levels of nutrients and trace metals (i.e., 4 µM phosphate (PO_4_^3-^), 60 µM nitrate (NO_3_^−^), 250 nM Fe, 50 nM Co, no Cr or Ni, (Sunda et al., 2005)), and 60 µM silicate (SiO_3_^2-^) was added to the ACM medium for culturing the two diatoms only.

For experiments, cultures were inoculated into the three Olivine Leachate (OL) treatments described below, with the addition of 4 µM phosphate (PO_4_^3-^) and 60 µM nitrate (NO_3_^−^). There was no nitrogen (N) added into the ACM or OL medium for the N_2_ fixers. Iron, Cobalt, Nickel (Fe, Co and Ni) and silicate (SiOH_4_) were not added to the OL medium, except as components of the olivine leachate (see below). The background nutrient concentrations in the collected natural seawater were 1µM NO_3_^−^, 0.1µM PO_4_^3-^ and 3µM SiOH_4_. Dissolved trace metals were not measured, but surface concentrations are typically very low (1nM Fe or less, (John et al., 2012)) at the SPOT time series site where the seawater was cleanly collected, relative to amounts added to the ACM and to the OL (**Table 1**) for phytoplankton culturing.

**Table 1.**
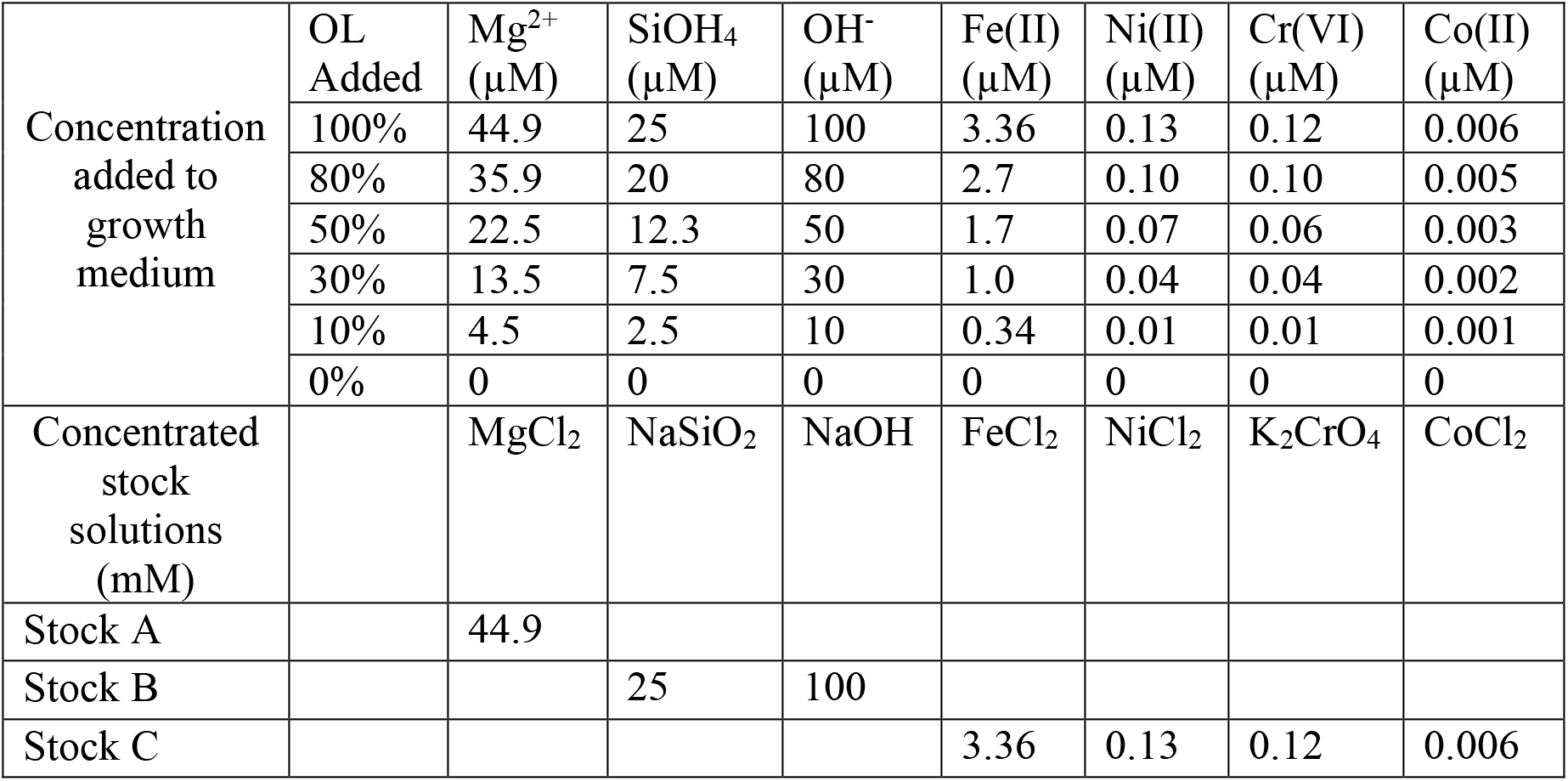
Concentrations of added ions or compounds in serial dilutions of synthetic olivine leachate (OL, 0% to 100%) used in the phytoplankton growth experiments; and concentrations of components in the three concentrated stocks used to prepare experimental medium (1mL/L added for 100% OL). Stock C was prepared in 10 nM HCl to keep the trace metals in solution until addition.

### Synthetic olivine leachate preparation

To simulate acute exposure of phytoplankton to elevated levels of olivine dissolution products in seawater, we prepared an artificial concentrated OL stock solution based on elemental analyses of commercial ground olivine rock (Sibelco. (2022) Technical Data - Olivine Refractory Grade Fine. Antwerp, Belgium). For experimental exposures, this concentrated OL stock was added to seawater growth medium to yield the final concentrations shown in **Table 1**, which will be referred to throughout as a “100%” concentration of OL. Experiments examining biological effects across a dilution range (0-100%) used correspondingly lower additions of the concentrated stock.

### Experimental methods

Semi-continuous culturing methods were used to achieve nearly steady-state growth. Cultures were diluted with fresh medium every 2 or 3 days, using in vivo fluorescence as a real time biomass indicator. Dilutions were calculated to bring the cultures back down to the biomass levels that were recorded after the previous day’s dilution. In this way, cultures were allowed to determine their own growth rates under each set of experimental conditions, without ever nearing stationary phase, significantly depleting nutrients or self-shading (Fu et al., 2022). For all experiments, cultures were sampled for a basic set of core biomass and physiological parameters, including cell counts, CO_2_ fixation, particulate organic carbon (POC), particulate organic nitrogen (PON), particulate organic phosphorus (POP) and biogenic silica (BSi, diatoms only) once steady-state growth was obtained for each growth condition (typically after 8−10 generations). Steady-state growth status was defined as no significant difference in cell- or in vivo-specific growth rates for at least 3 consecutive transfers.

#### There were four sets of experiments in this project

##### 1) Acute responses to elevated olivine leachate levels

The goal of this set of experiments was to investigate the responses of the diatoms *Nitzschia* and *Ditylum* to relatively high concentrations of olivine leachate, in order to determine acute exposure responses. To see if the leachate may have a positive or negative effect on their physiology, they were compared to their respective control cultures. There were a total of three treatments consisting of: OL (100%), ACM, and ACM with low Fe/Si (with 2 nM Fe EDTA added, and no added SiOH_4_).

##### 1. 2) Responses to a broad range of olivine leachate levels

In these experiments, *Synechococcus*, *Crocosphaera, Ditylum,* and *Emiliania huxleyi* were grown in culture medium across a series of OL dilutions (Table 1) to determine their responses across a range of leachate concentrations, from high to very low-level exposures.

##### 1. 3) Fe bioavailability and Cr toxicity from olivine leachate to N_2_-fixing cyanobacteria

The goal of this set of experiments was to investigate OL-derived Fe bioavailability to N_2_-fixing cyanobacteria, *Trichodesmium* and *Crocosphaera*. An additional experiment was conducted to investigate potential Cr(VI) toxicity.

##### 1. 4) Two species co-culture competition experiments during olivine leachate exposure

In order to test how OL may affect co-existence and competition between the diatom *Ditylum* and the coccolithophore *Emiliania huxleyi*, a simple batch co-culture competition experiment was carried out in which the 2 species were inoculated at a 1:1 ratio (based on equivalent levels of cellular Chlorophyll a due to the large differences in their cell sizes) into 100% OL and regular ACM, and grown for 10 days until early stationary phase. In vivo fluorescence and cell counts were monitored daily. Relative abundance and growth rates of the two species were determined based on microscopic cell counts during the exponential growth phase of the mixed cultures. Biogenic silica (BSi, an indicator of diatom abundance) and particulate inorganic carbon (PIC or calcite, an indicator of coccolithophore abundance) were collected every other day in order to further determine how these two species responded to co-culture with and without leachate additions.

### Analytical methods

#### Determination of growth rates and chlorophyll a

Growth rates were determined based on changes in chlorophyll a. For chlorophyll a determination, subsamples of 30 ml from each triplicate bottle were GF/F filtered, extracted in 6 ml of 90% acetone, stored overnight in the dark at −20°C, and chlorophyll a concentrations were measured fluorometrically using a Turner 10-AU fluorometer (Welschmeyer, 1994). Specific growth rates were determined using the equation:

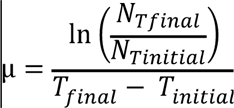

where μ is the specific growth rate (per day) and *N* is the chlorophyll a concentration at T_initial_ and T_final_ (Kling et al., 2021).

#### Particulate C, N, P and Si

Particulate organic carbon and nitrogen (CHN) samples from all experiments were filtered (pre-combusted GF/F) and frozen for analysis using a Costech Elemental Analyzer (Hutchins et al., 2007). Samples for biogenic silica (BSi) were filtered onto 25 mm diameter, 0.6 µm pore size polycarbonate filters (Pall Life Sciences), and analyzed according to (Brzezinski, 1985). POP (particulate organic phosphorus) samples were collected onto pre-combusted 25 mm GF/F filters and analyzed as in Fu et al. 2005 (Fu et al., 2005).

#### Primary productivity

For all species other than the coccolithophore (see below), primary production was measured in triplicate using 24h incubations (approximating net PP) with H^14^CO_3_ under the appropriate experimental growth conditions for each treatment (Fu et al., 2008). CO_2_ fixation rates were calculated using measured final experimental DIC concentrations and biomass. All samples for primary production were counted using a Wallac System 1400 liquid scintillation counter.

#### Photosynthetic and calcification rates of Emiliania huxleyi

For the coccolithophore, two 40 mL subsamples from each triplicate bottle were spiked with 0.5 µCi NaH^14^CO_3_. One subsample was incubated in the light and the other in the dark for 24 h. Then two sets of 20 mL aliquots from each sub-sample were filtered onto Whatman GF/F filters. The filters for photosynthetic rate determination were fumed with saturated HCl before adding scintillation cocktail fluid. Photosynthetic rate and calcification rate were calculated as described in Paasche et al. 1996 (Paasche et al., 1996).

#### Nitrogen fixation rates of Crocosphaera watsonii

In order to estimate the N_2_ fixation rates of *Crocosphaera*, initial and final particulate organic nitrogen samples (50mL) were collected on combusted GF/F filters over a 24 hr incubation. The PON specific N_2_ rates were calculated using the following equation:

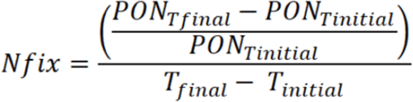

where Nfix is the N specific N_2_ fixation rates (day^−1^) and PON is the particulate organic nitrogen at Tinitial and Tfinal as measured using an elemental analyzer (Costech Analytical Technologies) (Fu et al., 2014)

#### Fe quota

Intracellular Fe content was determined by filtering culture samples onto acid-washed 0.2-μm polycarbonate filters (Millipore), and rinsing with oxalate reagent to remove extracellular trace metals (Tovar-Sanchez et al., 2003). Fe was determined with a magnetic sector-field high-resolution inductively coupled plasma mass spectrometer (ICPMS) (Element 2, Thermo) (Jiang et al., 2018; John et al., 2022).

#### Statistical methods

A one-way ANOVA analysis of variance was used to analyze differences between treatments using Prism 8. Differences between treatments were considered significant at p<0.05. Post-hoc comparisons were conducted using the Tukey’s multiple comparison test to determine any pairwise differences. Equality of variance was verified for all data using F tests, and the Shapiro Wilk test was used to test for significant departures from normality, which was not the case for any of our data sets. For experiments with only one or two OL treatments and the ACM control, graphs are presented with each treatment marked with a letter denoting significant differences at the p < 0.05 level from each of the other treatments. For experiments such as OL dilution series with many treatments (7 in this case), clear visualization of differences with all other treatments using letters is not feasible. For these experiments, significant differences in the OL treatments relative to the ACM positive control are indicated by asterisks (* = p < 0.05; ** = p < 0.01; *** = p < 0.001; **** = p < 0.0001). For all experiments, actual p values are given in the text.

## Results

### Diatoms

We hypothesized that the diatoms might benefit from the OL products Si and Fe, as they are both required for growth and can be limiting for this group (Tréguer et al., 2018). Hence, we grew the benthic diatom *Nitzschia* across three treatments: 100% OL alone, Aquil control medium (ACM), and ACM but with low, limiting Si and Fe concentrations (ACM-low-SF). *Nitzschia* grew and fixed carbon just as well in the 100% OL as the ACM (p=0.35; p=0.21), while showing reduced rates in the ACM-low-SF treatment (p=0.02; p< 0.0001; **Fig. 1A, B**). Likewise, the particulate Si:C ratios demonstrated 100% OL to be just as good a source of Si to *Nitzschia* as the ACM (p=0.98), and considerably better than the ACM-low-SF (p=0.0012; **Fig. 1C**).

**Fig 1.**
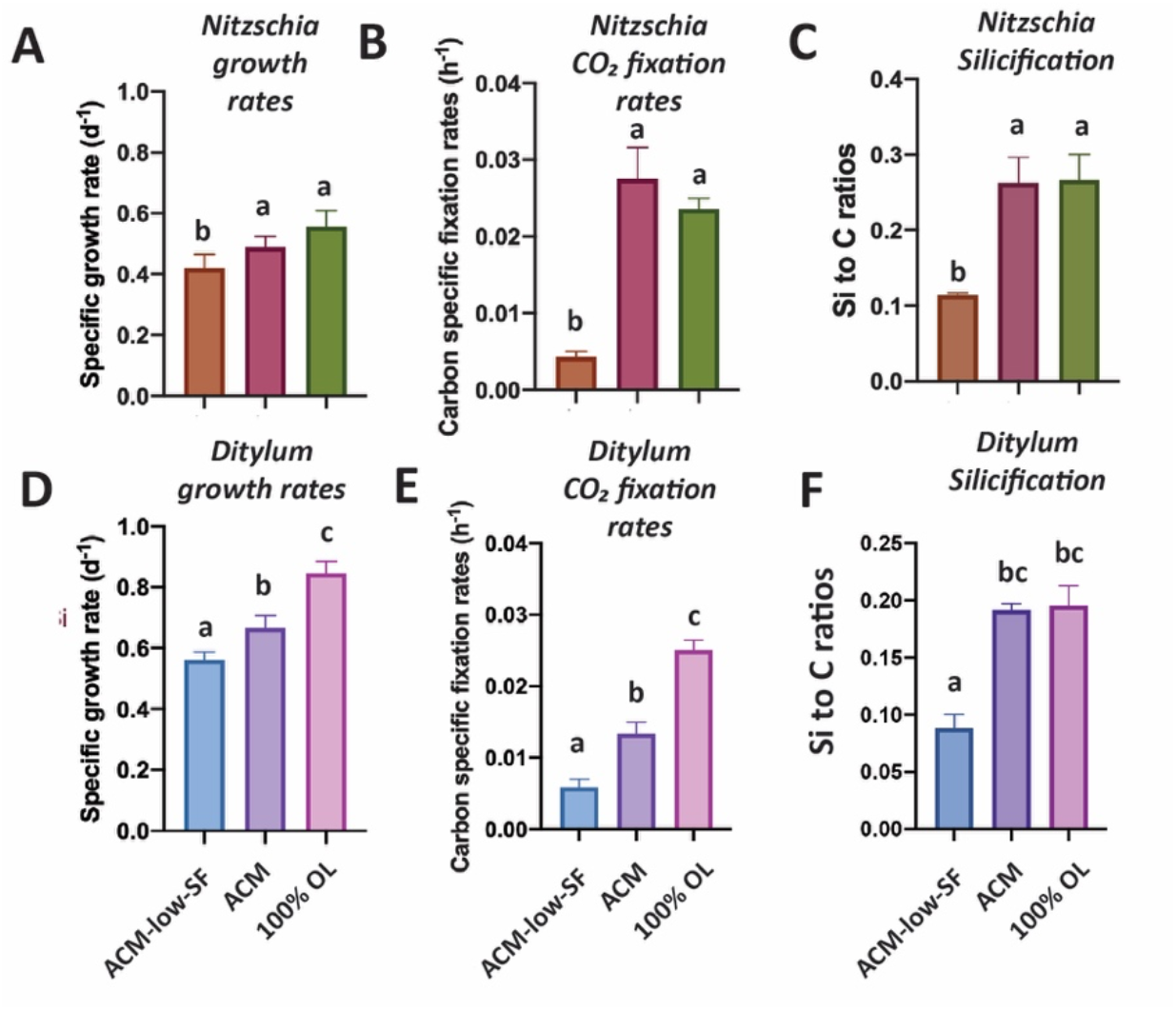
Effects of olivine leachate versus culture medium controls on growth and physiology of a benthic and pelagic diatom. **A) and D)** are Cell-specific growth rates (d^−1^), **B) and E)** are Carbon-specific fixation rates (hr^−1^) and **C) and F)** are Si:C ratios (mol:mol) for the diatoms *Nitzschia punctata* (benthic) and *Ditylum brightwellii* (pelagic), respectively. Abbreviations: OL is 100% olivine leachate, ACM is positive control Aquil medium, ACM-low-SF is positive control Aquil medium with lowered Si and Fe concentrations. Y-axis values represent the means, and error bars are the standard deviations of biological triplicate cultures for each treatment. Different letters indicate significant differences at the p < 0.05 level.

Growth and CO_2_ fixation rates of the planktonic diatom *Ditylum* were significantly higher in the 100% OL treatment compared to either the ACM (p=0.002; p=0.0001) or ACM-low-Si/Fe treatments (p=0.0002; p<0.0001; **Fig. 1D, E**), while Si:C ratios were the same (p=0.93; **Fig. 1F**). When *Ditylum* was grown across a range of OL concentrations (i.e., a dilution series from 0% to 100% additions, where 100% corresponds to the 100% OL treatment), we observed increasing growth and CO_2_ fixation rates with increasing OL concentrations, with maximum rates observed at and above 50% of the original OL that were not significantly different from those in the ACM control (p=0.06, 0.33, 0.99, **Fig. 2A, B)**. *Ditylum* particulate Si:C ratios also reached levels not significantly different to those seen in the ACM medium in the 100% additions (p=0.07, **Fig. 2C**). Likewise, *Ditylum* cellular Fe:P ratios measured by ICP-MS were not significantly different between 100% OL and ACM treatments, suggesting the diatom could access the same amount of Fe from the precipitated Fe(III) in the OL as from the soluble (EDTA-chelated) Fe(III) in the ACM culture medium (p=0.56; **Supp. Fig 1A**). These data demonstrate that even at extremely high concentrations, olivine dissolution products including trace metals were not toxic to these diatoms, but instead may provide sources of the essential nutrients iron and silicate to support their growth in nutrient replete conditions.

**Fig 2.**
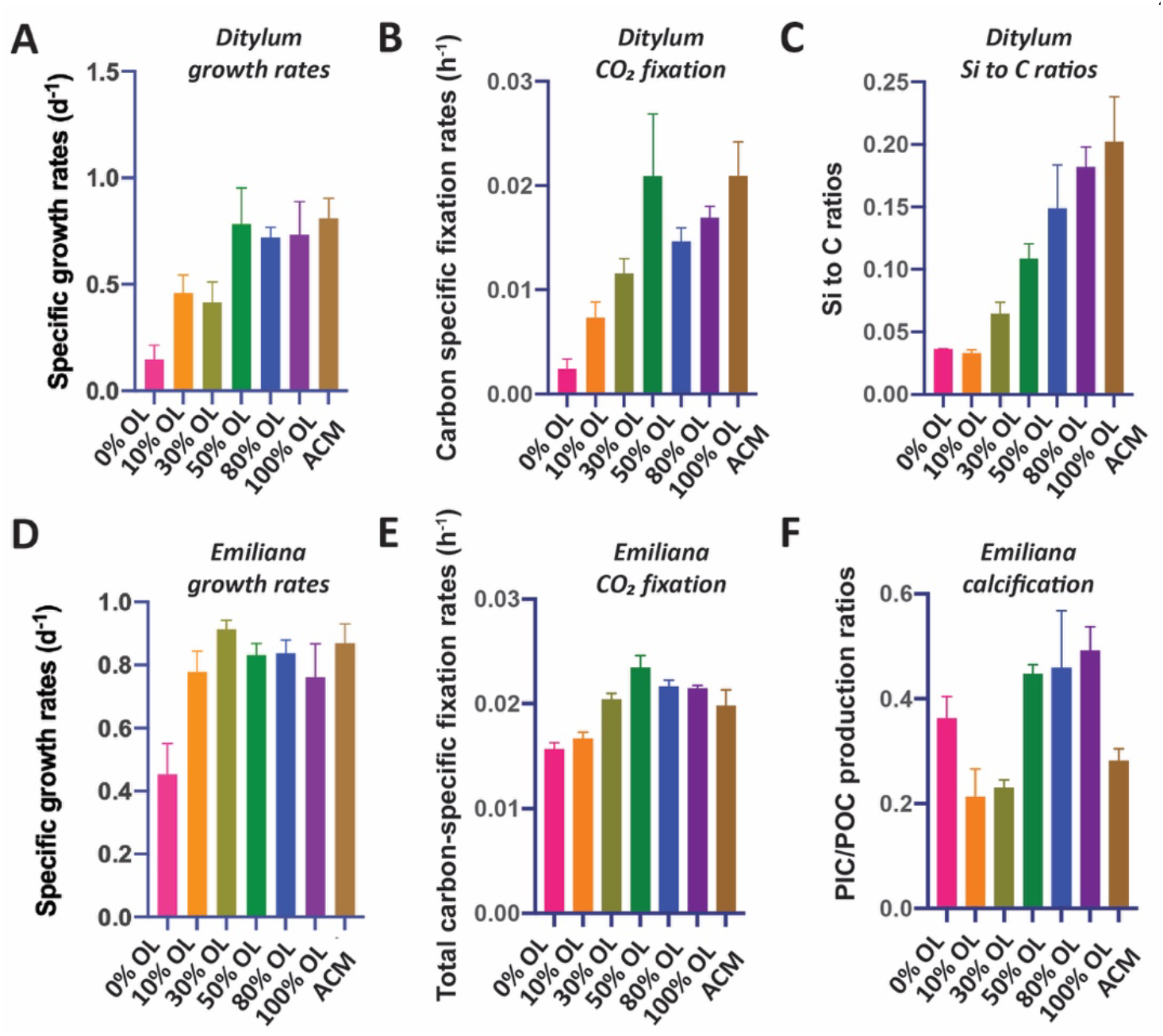
Effects of a dilution series of olivine leachate on growth and physiology of a marine diatom and coccolithophore. The diatom *Ditylum brightwellii* and the coccolithophore *Emiliana huxleyi* were grown across a range of dilutions of the olivine leachate (OL, 0-100%), and in the positive control medium (ACM). Shown are: **A)** Cell-specific growth rates (d^−1^), **B)** Carbon-specific fixation rates (hr^−1^) and **C)** and Si:C ratios (mol Silicon: mol Carbon) of *Ditylum brightwellii*, and: **D)** Cell-specific growth rates (d^−1^), **E)** Carbon-specific fixation rates (hr^−1^), and **F)** PIC/POC production ratios (calcification production rate/organic carbon fixation rate, unitless) of *Emiliana huxleyi* in the same OL and ACM treatments. Y-axis values represent the means and error bars are the standard deviations of biological triplicate cultures for each treatment. Relative to the ACM positive control, significant differences in OL treatments are indicated by * = p < 0.05; ** = p < 0.01; *** = p < 0.001; **** = p < 0.0001.

### Coccolithophores

It has been hypothesized that calcifying coccolithophores may benefit from an increase in alkalinity from olivine dissolution (Bach et al., 2019). In the OL dilution series, maximum growth rates for the coccolithophore *Emiliana* equivalent to those recorded in the ACM positive control medium were achieved at all added OL concentrations from 10% to 100% (p=0.36,0.92, 0.96, 0.98,0.26); the only significantly lower rate was at 0% OL (p<0.0001, **Fig. 2D**). POC production (CO_2_ fixation) rates were significantly reduced relative to the ACM in the 0% (p=0.0002) and 10% (p=0.002) treatments; but in all higher OL concentrations, primary production increased to levels that were the same as or higher than the ACM control (30% p=0.99, 50% p=0.04, 80% p=0.29, 100% p=0.45, 407 **Fig. 2E**).

Particulate inorganic carbon to particulate organic carbon production ratios (PIC:POC production ratios) were significantly higher at OL levels of 50-100% than in the ACM positive controls (p=0.009, 0.005, 0.001, **Fig. 2F**), possibly due to enhanced alkalinity in the high OL concentration treatments. PIC:POC production ratios were elevated in the 0% OL treatment relative to those in the 10% and 30% OL treatments due to CO_2_ fixation rates being reduced more than PIC fixation rates in this treatment, but in none of these treatments was this parameter significantly different from the ACM control (p=0.30,0.45,0.70, **Fig. 2F**).

An independent set of basic two-treatment experiments with the coccolithophore (ACM versus 100% OL, **Supp. Fig 2**) supported the results of the dilution series experiments shown in **Fig. 2**. *Emiliana* specific growth rates were slightly higher in the OL than in the ACM (p = 0.05), while cellular particulate inorganic:particulate organic carbon ratios (PIC:POC, mol:mol) were not significantly different in the two treatments (p= 0.08, **Supp. Fig. 2A**). Likewise, both POC-specific fixation rates (p=0.04; TC h^−1^) and PIC:POC production ratios were slightly higher in the OL than in the ACM treatments (p=0.05, **Supp. Fig. 2B**). Like the diatoms, these data demonstrate that olivine dissolution products are also not toxic to this common coccolithophore species, and that enhanced alkalinity may support marginally higher growth rates under nutrient replete conditions.

### Cyanobacteria

Like diatoms and coccolithophores, cyanobacteria could benefit from olivine dissolution due to their relatively high Fe (Hutchins and Boyd, 2016) and Ni requirements (Dupont et al., 2008). The OL dilution series experiments using the widely distributed picocyanobacterium *Synechococcus* showed positive responses in growth rates (**Fig. 3A**) and CO_2_ fixation rates (**Fig. 3B**) across the range of OL levels, similar to those of the eukaryotic algae. Both growth rates and carbon fixation rates were the same in the 100% OL treatment as in the ACM positive control treatment (p=0.94; p=0.46). ICP-MS measurements of *Synechococcus* cellular Fe:P ratios across a range of OL levels (0-100%) showed that this isolate accumulated much less Fe in the 0% OL than in the ACM treatment (p=0.02), but in all treatments with added OL, Fe:P ratios were the same as (10%. p=0.99, 30% 0.30, 50% 0.13, 100% 0.17) or higher than (80% p=0.01) than the ACM values (**Supp. Fig. 1B**). As with the eukaryotic phytoplankton tested, the synthetic OL provided a good source of Fe to support the growth of the picocyanobacterium.

**Fig 3.**
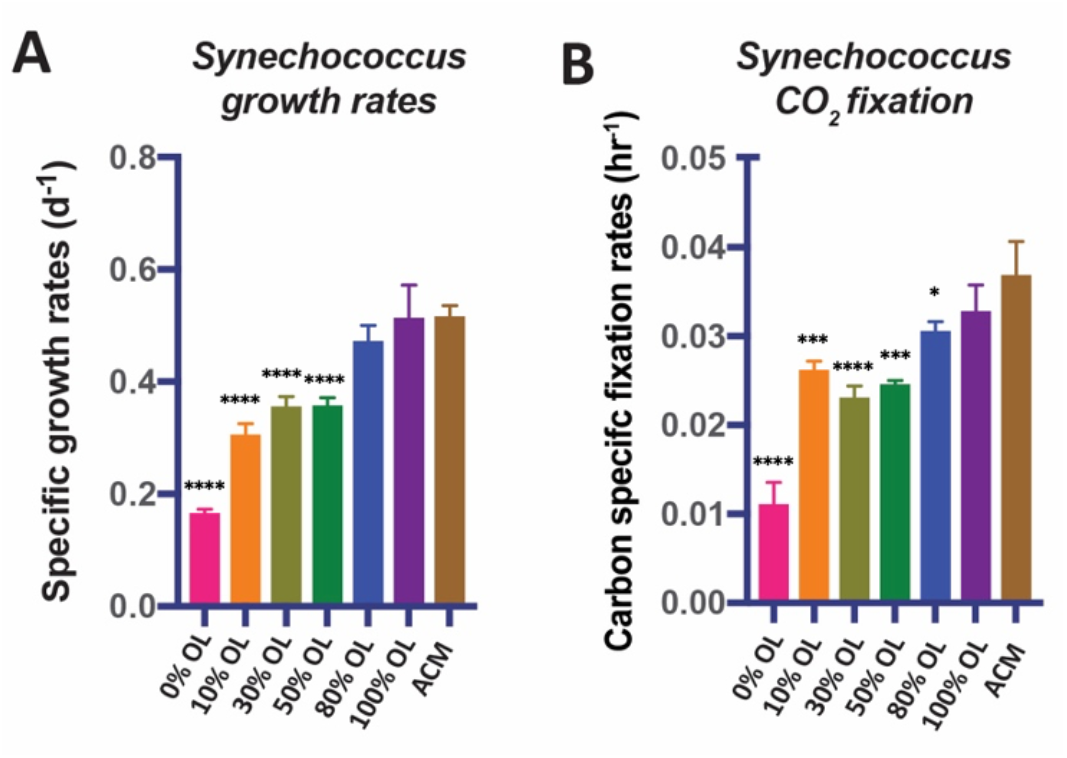
Effects of a dilution series of olivine leachate on growth and CO_2_ fixation of a marine cyanobacterium. The unicellular picocyanobacterium *Synechococcus* sp. was grown across a range of dilutions of the olivine leachate (OL, 0-100%), and in the positive control medium (ACM). Shown are: **A)** Cell-specific growth rates (d^−1^) and **B)** Carbon-specific fixation rates (hr^−1^). Values represent the means and error bars are the standard deviations of triplicate cultures for each treatment. Relative to the ACM positive control, significant differences in OL treatments are indicated by * = p < 0.05; ** = p < 0.01; *** = p < 0.001; **** = p < 0.0001.

In striking contrast to the eukaryotic algae and the non-diazotrophic (i.e., non-N_2_-fixing) picocyanobacterium *Synechococcus*, the N_2_-fixing cyanobacterium *Trichodesmium* could not grow at any concentration of OL tested **(Fig. 4A)**. One possible explanation for this lack of growth is toxic effects by one of the trace metal components of the OL. We hypothesized that added levels of Ni and Co are unlikely to be toxic, as these nutrient metals have been found to be relatively non-toxic to many phytoplankton at similar environmental concentrations (Guo et al., 2022; Karthikeyan et al., 2019; Panneerselvam et al., 2018). Hence, we hypothesized that Cr toxicity should be considered as a likely possible scenario (Frey et al., 1983; Kiran et al., 2016). Another possibility is that *Trichodesmium* did not experience toxic effects but instead was unable to access Fe from OL. This N_2_-fixer requires more Fe than virtually any other phytoplankton species (Hutchins and Sañudo-Wilhelmy, 2021), and the OL was the only source of Fe provided in our experiments. Fe(II) released into seawater from olivine dissolution likely quickly oxidizes to Fe(III), which then precipitates and becomes insoluble at the elevated concentrations in our OL (Manck et al., 2022). This could render it biologically unavailable to the cellular Fe uptake systems of some species. We deliberately designed our OL to replicate this oxidation/precipitation process, and as expected observed visible reddish-brown amorphous colloidal Fe precipitates on the bottom of the growth flasks for all synthetic OL treatments.

**Fig 4.**
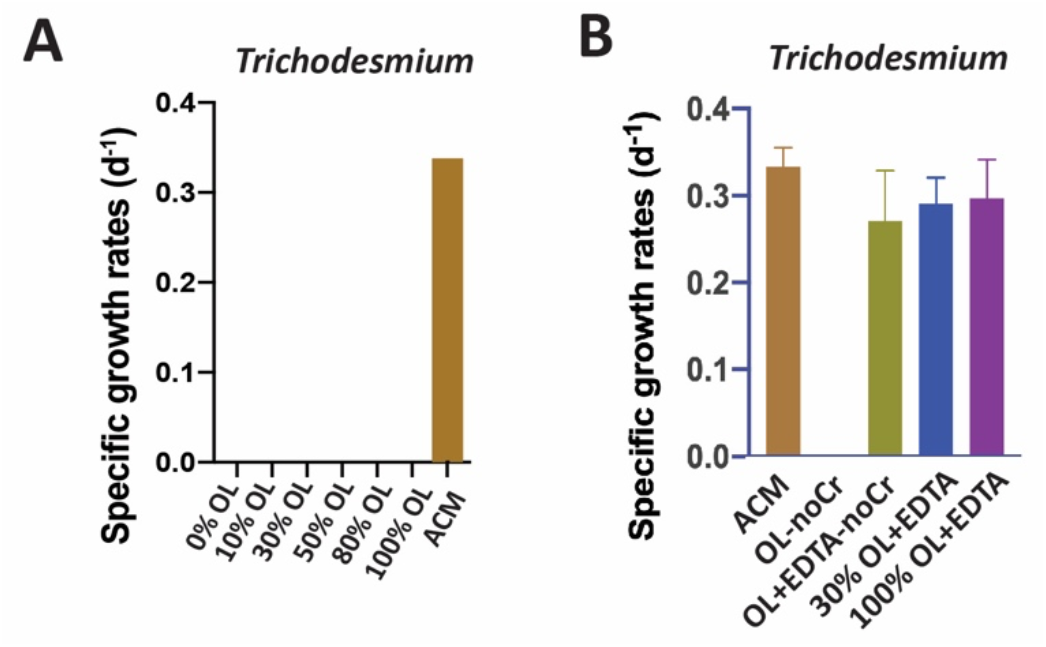
Effects of olivine leachate versus culture medium controls on growth of a colonial marine N_2_-fixing cyanobacterium. Shown are **A)** Cell-specific growth rates (d^−1^) of the colonial cyanobacterium *Trichodesmium erythraeum* across a range of dilutions of the olivine leachate (OL, 0-100%) and in the positive control medium (ACM), and **B)** Cell-specific growth rates (d^−1^) of *Trichodesmium* in two concentrations of OL (30% and 100%) with the synthetic metal chelator EDTA, and in OL without Cr or EDTA, and in OL without Cr but plus EDTA, all versus ACM. Unless growth rates were zero, relative to the ACM positive control, significant differences in OL treatments are indicated by * = p < 0.05; ** = p < 0.01; *** = p < 0.001; **** = p < 0.0001.

Accordingly, we designed another set of experiments to test for both lack of Fe bioavailability and specific sensitivity to Cr, as has been done in previous cyanobacterial studies (Kiran et al., 2016). To do this, we formulated several variants of the olivine leachate: 1) normal OL (100% concentration), 2) OL (100% concentration) with a synthetic ligand (EDTA) that solubilizes Fe(III), and thus makes it broadly bioavailable (OL+EDTA), 3) OL (100% concentration) but with no Cr (OL-noCr), 4) and OL (100% concentration) but with no Cr and with EDTA (OL+EDTA-noCr) **(Fig. 4B**). *Trichodesmium* also could not grow in the OL medium without added Cr (OL-noCR, **Fig. 4B)**, demonstrating that the lack of growth observed in OL was not due to Cr toxicity.

However, growth recovered to the same levels as in the ACM in all three treatments where EDTA was added (30%OL+EDTA p=0.62, 100%OL+EDTA p=0.84, 100%OL+EDTA-noCr p=0.30) to the leachate **(Fig. 4B)**. Together, these results suggest that poor bioavailability of the precipitated Fe(III) (and not Cr toxicity) was the likely cause for *Trichodesmium’*s inability to grow in the unmodified OL.

OL also reduced the growth rates of the unicellular N_2_-fixing cyanobacterium *Crocosphaera*, although not to the same extent as for *Trichodesmium,* which didn’t grow at all without the addition of EDTA*. Crocosphaera* exhibited no growth at 0% OL, likely due to severe Fe limitation. From 10% to 100% OL, growth rates were 22-44% of those in ACM (p<0.001), and growth partially recovered in 100% OL+EDTA to 76% of rates in ACM (p<0.0001, **Fig. 5A**). Results were very similar for CO_2_ fixation rates and N_2_-fixation rates in OL, which were severely reduced by 64-100% (carbon fixation, p<0.0001) and 69-88% (N_2_ fixation, p<0.0001) relative to ACM in all OL treatments, but reached maximum values of 80% (p=0.002) and 63% (p<0.0001) of ACM treatment rates, respectively, when EDTA was added to the OL (**Fig. 5B,C**). This suggests that oxidized Fe from OL was not effectively utilized to support growth for either of the two N_2_-fixing cyanobacteria tested, in contrast to the diatoms, coccolithophores, and *Synechococcus*. Their growth recovery after EDTA additions indicates that the other trace metals Cr, Ni, and Co in the olivine leachate were likely not toxic, even at extremely high concentrations. Interestingly, unlike *Trichodesmium* which could not grow at all on OL alone but recovered fully upon EDTA additions, *Crocosphaera* could still grow at lower rates on OL but could not grow as fast upon EDTA additions as in ACM. Future experiments are needed to understand these differences in species-specific responses between these two N_2_-fixers. Taken together, these data suggest that when olivine dissolves in seawater, it is unlikely to provide a readily bioavailable Fe source to diazotrophic cyanobacteria, although this does not preclude them obtaining Fe from their usual natural sources such as other sediments, rivers, dust inputs etc. Thus, it seems likely that olivine may have negligible or no effect (positive or negative) on the physiology of these cyanobacteria, although further work will be needed to put these results into a more realistic ecological context to understand the full responses of N_2_ fixing cyanobacteria to olivine dissolution.

**Fig 5.**
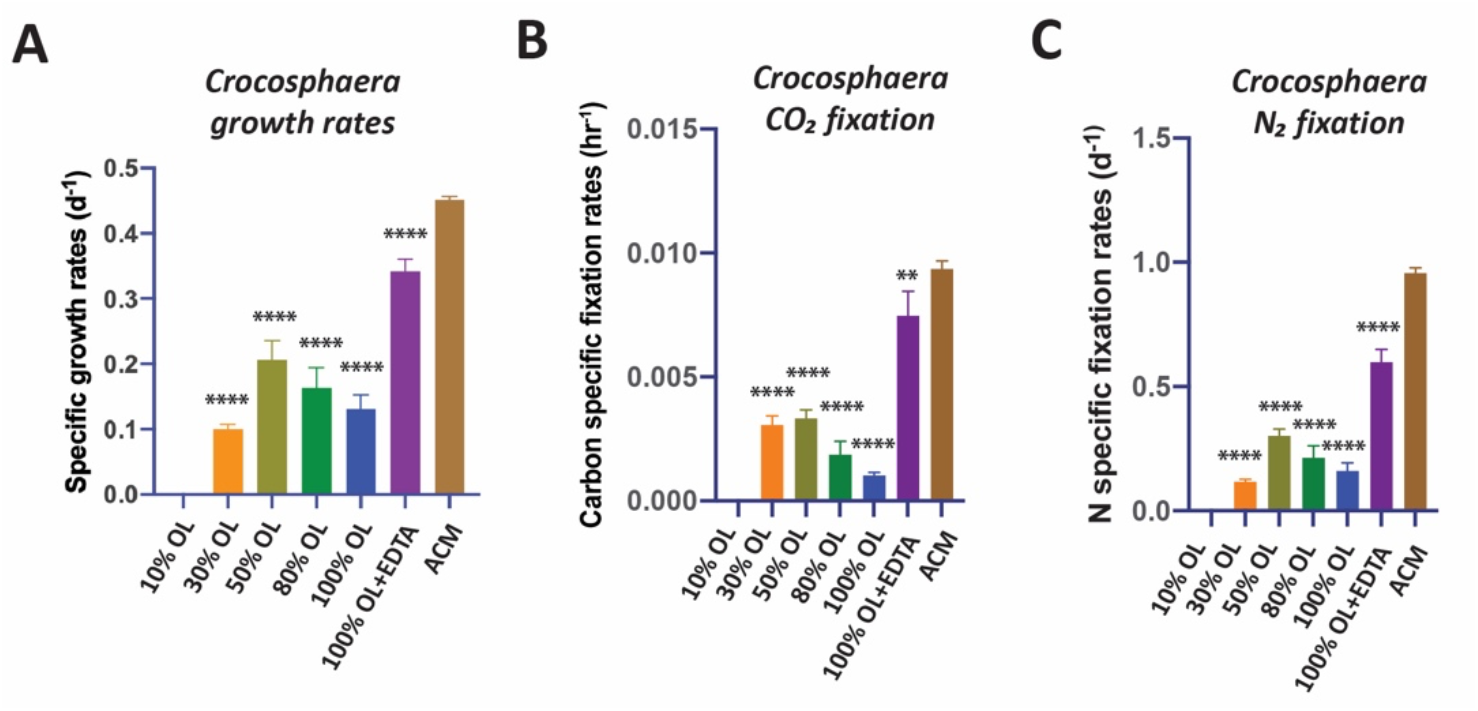
Effects of olivine leachate dilution series versus culture medium controls on the physiology of a unicellular marine N_2_-fixing cyanobacterium. **A)** Cell-specific growth rates (d^−1^), **B)** Carbon-specific fixation rates (hr^−1^), and **C)** N-specific fixation rates (day^−1^) of the unicellular cyanobacterium *Crocosphaera watsonii* grown across a range of dilutions of the olivine leachate (OL, 0-100%), in 100% OL plus EDTA (OL+EDTA), and in the positive control medium (ACM). Values represent the means and error bars are the standard deviations of triplicate cultures for each treatment. Relative to the ACM positive control, significant differences in OL treatments are indicated by * = p < 0.05; ** = p < 0.01; *** = p < 0.001; **** = p < 0.0001.

### Diatom/Coccolithophore Competitive Co-culture

Results of the co-culture, or competition, experiment with the diatom *Ditylum brightwellii* and the coccolithophore *Emiliania huxleyi* are shown in **Fig. 6**. Unlike the semi-continuous experiments shown in the previous figures, this experiment used closed-system “batch” culturing methods in order to assess and compare effects on relative biomass accumulation by each species over time. OL (100% concentration) supported growth of both the diatom (**Fig. 6A**) and the coccolithophore (**Fig. 6B**) in mixed culture, and biomass was very similar for both species between the OL and ACM treatments throughout most of the experiment. However, cell yields were higher in the ACM at the final timepoint for the diatom (p = 0.009, **Fig. 6A**). Final cell counts were also higher in the ACM for the coccolithophore, but this difference was not significant (p= 0.31; **Fig. 6B**). Similar trends were observed when diatom biomass was estimated as biogenic silica (BSi, **Fig. 6C**, p =0.002) and when coccolithophore biomass was assessed as calcite or particulate inorganic carbon (PIC, **Fig. 6D**, p = 0.04). For the diatom, OL supported growth rates similar to those in the ACM treatment during the first half of the experiment (**Fig. 6A,C**; **Supp. Fig. 3A**). Growth rates were also similar in the OL and ACM mediums for the coccolithophore (**Fig. 6B,D; Supp. Fig. 3B**). Hence, both phytoplankton species were able to grow similarly well in co-culture, where neither exhibited any strong competitive advantage over the other.

**Fig. 6.**
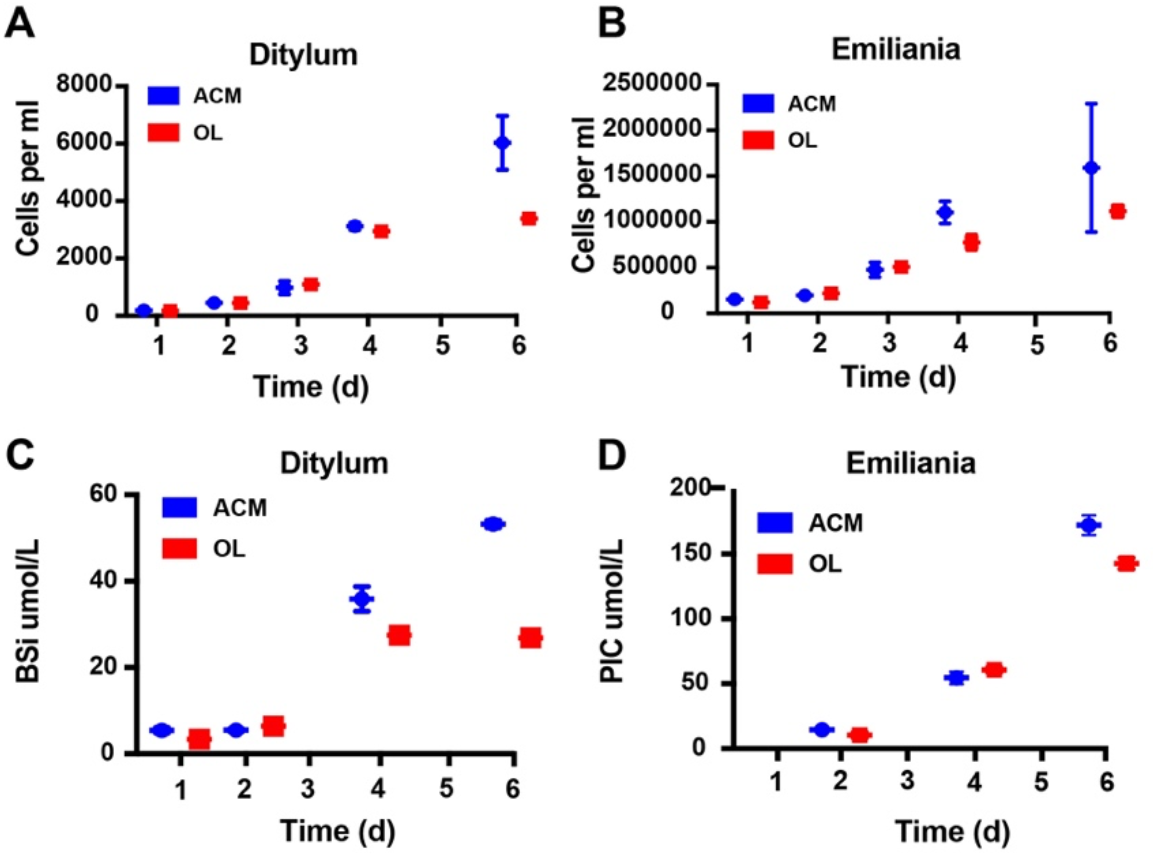
Effects of olivine leachate versus culture medium controls on growth competition and biomineralization during co-culture of a diatom and a coccolithophore. Shown are 5-day growth curves (cells mL^−1^) for **A)** the diatom *Ditylum brightwellii* and **B)** the coccolithophore *Emiliana huxleyi* in mixed cultures grown in olivine leachate (OL, red symbols) and positive control medium (ACM, blue symbols). Also shown are **C)** Biogenic silica (BSi, μmol L^−1^), a proxy for diatom biomass, and **D)** Calcite or particulate inorganic carbon (PIC, μmol L^−1^), a proxy for coccolithophore biomass, in the OL and ACM treatments in the same growth competition experiment. Values represent the means and error bars are the standard deviations of triplicate cultures for each treatment.

## Discussion

In general, we observed no toxic effects from our simulated olivine dissolution products even at extremely high concentrations across all the phytoplankton species tested, consistent with other recent observations examining OAE scenarios (Gately et al., 2023) and trace metals (Guo et al., 2022). Guo et al. (2022) particularly focused on exposing 11 phytoplankton groups to elevated Ni concentrations and did not observe strong effects across these taxa. Although it is unknown what chemical species of dissolved Ni primarily influence phytoplankton physiology, most studies indicate that phytoplankton primarily interact with free Ni^2+^ ions but are not particularly sensitive to the total dissolved Ni concentration (Guo et al., 2022). Guo et al. (2022) and our study used the same Ni-containing compound, NiCl, as a source of Ni^2+^. Guo et al. also used the same base Aquil control medium as our ACM. ACM contains EDTA that binds with metal ions like Ni to improve their dissolution, which subsequently lowers the free Ni ion concentrations (e.g., Ni^2+^) relative to the total dissolved Ni pool. Although broad negative effects of enhanced Ni concentrations were not observed across taxa, Guo observed some species-specific responses across variations in (0-100 µM) EDTA and Ni (0-50 µM) concentrations, indicating that specific phytoplankton groups are impacted differently depending on the chemical species in the total dissolved Ni pools and/or the concentration and type of organic ligands in seawater. Our results are generally consistent with their overall findings, as the phytoplankton groups tested here did not exhibit negative effects upon elevated exposure to Ni with (e.g., ACM) and without added EDTA (e.g., OL), suggesting that Ni was not toxic irrespective of the concentration of different Ni species in the dissolved pool or that of the Ni^2+^ ion. However, our experiments were not designed to test for taxon-specific differences in responses to specific Ni species or variations in EDTA concentrations.

The two diatoms were able to use synthetic OL-derived Si and Fe to support near-maximum growth rates and carbon fixation rates, as well as robust silica frustule development (as assessed by cellular Si:C ratios); both of these nutrients can frequently limit diatom growth in various parts of the ocean (Tréguer et al., 2018; Hutchins and Boyd, 2016). OL-derived alkalinity and iron increases also supported growth of the tested coccolithophore, consistent with previous observations showing increases in red algae calcification (Gore et al., 2019) and net reef calcification calcifying corals (Albright et al., 2016) in response to OAE. Similarly, a representative of the globally distributed, picocyanobacterium *Synechococcus* increased its growth and carbon fixation rates as OL concentrations increased. Although OL could not support *Trichodesmium* and *Crocosphaera* growth due to their inability to use Fe(III), olivine dissolution products were not observed to be toxic. Their inability to use Fe(III) is a neutral effect due to other sources of bioavailable Fe in the water column (Hutchins and Boyd, 2016).

Thus, these results using 6 model species suggest that many phytoplankton species may not be negatively impacted by even high levels of elements derived from olivine dissolution, and that some olivine dissolution products may support their growth, primary productivity, and biomineralization when OL is available at high enough concentrations in certain environmental conditions. For example, it is important to note that potential growth benefits to phytoplankton *in situ* will also depend on ambient concentrations of other important nutrients, such as nitrogen (N) and phosphorus (P). Our cultures contained an abundance of other required nutrients, thus enabling phytoplankton to take advantage of dissolution products for growth (e.g., Si, Fe, alkalinity). However, if nutrients like N and P are primarily limiting in natural environments, then olivine dissolution products are not expected to have any growth effect. In addition, these cultures represent closed systems that do not allow olivine products to be diluted with fresh seawater. In natural settings, advection in both sediment porewaters (Reimers et al., 2004) and the water column (He and Tyka, 2022) will lead to short residence times, thereby rapidly diluting olivine dissolution products. Hence, these physical dynamics will prevent high concentrations of olivine dissolution products from accumulating in seawater in coastal systems. Thus, even the most dilute leachate treatment in this study is likely more concentrated than the anticipated concentrations of olivine dissolution products expected under field conditions. It is important to note though that our experiments focused on physiological responses, while further work will be needed to explore the possibility of indirect effects on important ecological factors such as predation or competition. Further research will also be needed to test for direct and indirect effects using other species of phytoplankton not examined here and belonging to other important functional groups.

Bach et al. (2019) (Bach et al., 2019) hypothesized that under nutrient-limited conditions, silicate, iron and nickel releases from marine applications of silicate minerals like olivine might particularly benefit diatoms and cyanobacteria, as these groups have especially high requirements for one or more of these nutrients. Thus, they expected that olivine applications might produce a “Greener” ocean. They also suggested that adding minerals derived from CaCO_3_^−^ (such as quicklime applications) would particularly favor coccolithophores, due to rapidly enhanced seawater alkalinity. This outcome would produce a “Whiter” ocean (the color of coccolithophore calcite). Although we did not test CaCO_3_^−^ derivatives, our results with nutrient-replete synthetic OL seem to represent a “Green and White” ocean scenario, since in individual experiments diatoms, picocyanobacteria, and coccolithophores all responded positively to OL at the relatively elevated levels applied in our experiments. This conclusion is further supported by the results of our nutrient replete diatom/coccolithophore co-culture experiment, which showed that OL stimulated both species simultaneously rather than conferring a competitive advantage on one or the other. This suggests that in the ocean, competitive outcomes between diatoms and coccolithophores may not be affected by olivine dissolution under nutrient-replete conditions, although we did not test this under nutrient-limited conditions that are commonly encountered by phytoplankton in nature.

Iron in olivine minerals is present as reduced Fe(II), and we added it in this form to our synthetic OL. However, when Fe(II) dissolves in oxic seawater, it quickly (within minutes) oxidizes to highly insoluble Fe(III), which precipitates out as amorphous iron hydroxides (Millero et al., 1987). Clearly, in our experiments this oxidized particulate iron must have been available to the species that showed growth enhancement with OL, since no other iron source was provided in the seawater growth medium. In accordance with their well-studied reductive Fe uptake systems (Morrissey and Bowler, 2012), there is evidence that diatoms can access Fe to some degree from freshly precipitated amorphous colloidal Fe hydroxides (like those in our experiments), although the bioavailability of Fe precipitates declines quickly as the hydroxides age and acquire a more crystalline structure (Yoshida et al., 2006). Alternately, the precipitated Fe in our experiments could have become available to the phytoplankton ferric reductases via solubilization by siderophores produced by bacteria in our non-axenic cultures (Coale et al., 2019). In some cases diatoms, can also potentially take up Fe through endocytosis (Kazamia et al., 2018).

The responses of the two N_2_-fixing cyanobacteria were in striking contrast to those of the other three phytoplankton groups tested. These diazotrophs were either unable to grow in our artificial OL at all (*Trichodesmium*), or could only grow to a very limited degree (*Crocosphaera*). However, our results with experimental additions of the artificial iron chelator EDTA (ethylene diamine tetra acetic acid) suggest that other mechanisms may enable iron bioavailability. For example, previous research has suggested that *Trichodesmium* cannot directly access particulate Fe(III) forms, but likely relies on bacteria residing on and in natural colonies to produce siderophores, which then solubilize particulate Fe(III) sources and make them bioavailable (Rubin et al., 2011; Lee et al., 2018). Since cultured *Trichodesmium* such as ours typically do not produce colonies, but grow instead as individual filaments of cells, cultures of this diazotroph are likely deficient in many of these iron-acquiring microbial symbionts (Rubin et al., 2011). The iron uptake systems of *Crocosphaera* have been less well-characterized, but like *Trichodesmium*, molecular studies suggest this unicellular diazotroph lacks the genetic capacity to produce endogenous siderophores (Shi et al., 2010; Yang et al., 2022). Our results show that when we add the artificial iron chelator EDTA (which substitutes for ligands produced by the missing bacteria in cultures), the synthetic OL supports near-maximum growth of both diazotrophs. Thus, reduced growth rates of these cyanobacteria in OL without EDTA appear to be due to severe iron limitation, not toxicity of any OL component. In our experiments the cells were forced to grow on OL as a sole source of iron, but in coastal ecosystems where olivine deployments would occur, there are typically many other natural sources of iron to support algal growth (Capone and Hutchins, 2013; Hutchins and Boyd, 2016). In nature, *Trichodesmium* is also likely to occur mostly as colonies, and so may have access to additional siderophore-bound iron, including from both naturally occurring supplies as well as potentially from any oceanic olivine applications. Thus, changes in the iron nutritional status of N_2_ fixers due to olivine additions in-situ may not occur in the real ocean.

While the reduced growth rates of the diazotrophs on our synthetic OL appears to be due to iron limitation, our experiments also shed light on potential effects of other trace metals present in the formulation. Of the metals found in our synthetic OL, Ni and Co are considered nutrient elements with relatively low toxicity; in fact, the concentrations added even in our maximum dosage experiments were well below those that have been reported to be toxic to phytoplankton (Karthikeyan et al., 2019; Vink and Knops, 2023). However, Cr has the potential to be biologically problematic. Cr(III) found in olivine is relatively insoluble, so in this form it is probably not a major source of exposure for planktonic organisms. However, if it oxidizes to Cr(VI), it becomes much more soluble, and thus more bioavailable and potentially toxic. Cr(III) oxidation is thermodynamically unfavorable, but can be facilitated by borate ions always present in seawater, or by the presence of biologically or photochemically-produced oxidants like H_2_O_2_ (Pettine et al., 1991), and by naturally occurring manganese oxides (Weijden and Reith, 1982). For these reasons, following the principle of “worst case scenario”, we used a soluble Cr(VI) salt in our synthetic OL formulation. Despite this, we found that the presence or absence of the relatively elevated levels of dissolved Cr(VI) in our regular synthetic OL did not make any difference to the growth of *Trichodesmium* or the other tested phytoplankton species. Particularly, because synthetic OL stimulated near-maximum growth rates in the diatoms, coccolithophore and picocyanobacterium, we presume that the Cr(VI) additions did not adversely affect these groups either.

The goal of this work was to test both extreme levels and simultaneous exposure of multiple, biologically important olivine dissolution products that could influence microbial physiology in order to identify thresholds and response curves. Accordingly, our experiments focused on determining acute effects of high concentrations of olivine dissolution products. In general, they suggest that negative impacts may be few even for large olivine deployments, given the high concentrations of tested olivine dissolution products. Because these microplankton serve as important links to higher trophic levels, these data suggest minimal long-term impacts from olivine dissolution on ecosystem services. Future research directions may include longer term experiments with prokaryotes and natural microbial communities to expand our understanding of olivine exposure on important taxa that help drive biogeochemical cycling in the oceans, particularly experiments to test for ecological effects on processes like competition and trophic interactions at the community and ecosystem levels. Similar experiments can also be conducted except with other OAE feedstocks harboring different chemical compositions and more rapid dissolution timescales (Renforth and Henderson, 2017). Future studies can also focus on determining how biological processes like photosynthesis, respiration, and organic ligand production could influence olivine dissolution kinetics and their impacts on carbon dioxide removal.

## Data Availability

All data can be found at https://zenodo.org/record/8157750.

## Author contribution

D.A.H., F.-X.F., S.J.R., and N.G.W. designed the research; D.A.H., F.-X.F., S.-C.Y, and S.G.J. performed the research. D.A.H., F.-X.F., S.-C.Y, N.G.W., and S.G.J. analyzed the data. D.A.H., F.-X.F., S.-C.Y, N.G.W., S.J.R., M.G.A., and S.G.J. wrote the paper.

## Competing interests

Authors D.A.H. and F.-X.F. received research funding from Vesta, PBC. N.G.W., M.G.A., and S.J.R. are full time employees at Vesta, PBC.

## Notes

https://zenodo.org/record/8157750

### Summary of Updates

This revision has updated text and framing after 2 reviewer comments as a result of submitting the manuscript for review here (https://egusphere.copernicus.org/preprints/2023/egusphere-2023-930/). Please see the open discussion for comments and responses via the URL.

## References

Albright, R., Caldeira, L., Hosfelt, J., Kwiatkowski, L., Maclaren, J. K., Mason, B. M., Nebuchina, Y., Ninokawa, A., Pongratz, J., Ricke, K. L., Rivlin, T., Schneider, K., Sesboüé, M., Shamberger, K., Silverman, J., Wolfe, K., Zhu, K., and Caldeira, K.: Reversal of ocean acidification enhances net coral reef calcification, Nature, 531, 362–365, https://doi.org/10.1038/nature17155, 2016.

Bach, L. T., Gill, S. J., Rickaby, R. E. M., Gore, S., and Renforth, P.: CO2 Removal With Enhanced Weathering and Ocean Alkalinity Enhancement: Potential Risks and Co-benefits for Marine Pelagic Ecosystems, Frontiers in Climate, 1–21, https://doi.org/10.3389/fclim.2019.00007, 2019.

Beerling, D. J., Kantzas, E. P., Lomas, M. R., Wade, P., Eufrasio, R. M., Renforth, P., Sarkar, B., Andrews, M. G., James, R. H., Pearce, C. R., Mercure, J.-F., Pollitt, H., Holden, P. B., Edwards, N. R., Khanna, M., Koh, L., Quegan, S., Pidgeon, N. F., Janssens, I. A., Hansen, J., and Banwart, S. A.: Potential for large-scale CO2 removal via enhanced rock weathering with croplands, Nature, 1–20, https://doi.org/10.1038/s41586-020-2448-9, 2021.

Bertrand, E. M., Allen, A. E., Dupont, C. L., Norden-Krichmar, T. M., Bai, J., Valas, R., and Saito, M. A.: Influence of cobalamin scarcity on diatom molecular physiology and identification of a cobalamin acquisition protein, Proceedings of the National Academy of Sciences, E1762– E1771, https://doi.org/10.1073/pnas.1201731109/-/dcsupplemental, 2012.

Brzezinski, M. A.: THE Si:C:N RATIO OF MARINE DIATOMS: INTERSPECIFIC VARIABILITY AND THE EFFECT OF SOME ENVIRONMENTAL VARIABLES, J Phycol, 21, 347–357, https://doi.org/10.1111/j.0022-3646.1985.00347.x, 1985.

Capone, D. G. and Hutchins, D. A.: Microbial biogeochemistry of coastal upwelling regimes in a changing ocean, Nat Geosci, 6, 711–717, https://doi.org/10.1038/ngeo1916, 2013.

Caserini, S., Storni, N., and Grosso, M.: The Availability of Limestone and Other Raw Materials for Ocean Alkalinity Enhancement, Glob. Biogeochem. Cycles, 36, https://doi.org/10.1029/2021gb007246, 2022.

Coale, T. H., Moosburner, M., Horák, A., Oborník, M., Barbeau, K. A., and Allen, A. E.: Reduction-dependent siderophore assimilation in a model pennate diatom, Proceedings of the National Academy of Sciences, 1–9, https://doi.org/10.1073/pnas.1907234116, 2019.

Dupont, C. L., Barbeau, K., and Palenik, B.: Ni Uptake and Limitation in Marine *Synechococcus* Strains, Appl Environ Microb, 74, 23–31, https://doi.org/10.1128/aem.01007-07, 2008.

Field, C., Behrenfeld, M., Randerson, J., and Falkowski, P.: Primary production of the biosphere: integrating terrestrial and oceanic components, Science, 281, 237–240, 1998.

Flipkens, G., Blust, R., and Town, R. M.: Deriving Nickel (Ni(II)) and Chromium (Cr(III)) Based Environmentally Safe Olivine Guidelines for Coastal Enhanced Silicate Weathering, Environ Sci Technol, 55, 12362–12371, https://doi.org/10.1021/acs.est.1c02974, 2021.

Frey, B. E., Riedel, G. F., Bass, A. E., and Small, L. F.: Sensitivity of estuarine phytoplankton to hexavalent chromium, Estuar Coast Shelf Sci, 17, 181–187, https://doi.org/10.1016/0272-7714(83)90062-8, 1983.

Fu, F., Zhang, Y., Bell, P. R. F., and Hutchins, D. A.: PHOSPHATE UPTAKE AND GROWTH KINETICS OF *TRICHODESMIUM* (CYANOBACTERIA) ISOLATES FROM THE NORTH ATLANTIC OCEAN AND THE GREAT BARRIER REEF, AUSTRALIA1, J Phycol, 41, 62–73, https://doi.org/10.1111/j.1529-8817.2005.04063.x, 2005.

Fu, F., Tschitschko, B., Hutchins, D. A., Larsson, M. E., Baker, K. G., McInnes, A., Kahlke, T., Verma, A., Murray, S. A., and Doblin, M. A.: Temperature variability interacts with mean temperature to influence the predictability of microbial phenotypes, Global Change Biol, 28, 5741–5754, https://doi.org/10.1111/gcb.16330, 2022.

Fu, F. X., Yu, E., Garcia, N. S., Gale, J., Luo, Y., Webb, E. A., and Hutchins, D. A.: Differing responses of marine N2 fixers to warming and consequences for future diazotroph community structure, Aquatic Microbial Ecology, 72, 33–46, https://doi.org/10.3354/ame01683, 2014.

Fu, F.-X., Mulholland, M. R., Garcia, N. S., Beck, A., Bernhardt, P. W., Warner, M. E., Sañudo-Wilhelmy, S. A., and Hutchins, D. A.: Interactions between changing pCO_2_, N_2_ fixation, and Fe limitation in the marine unicellular cyanobacterium *Crocosphaera*, Limnology and Oceanography, 53, 2472–2484, https://doi.org/10.4319/lo.2008.53.6.2472, 2008.

Gately, J. A., Kim, S. M., Jin, B., Brzezinski, M. A., and Iglesias-Rodriguez, M. D.: Coccolithophores and diatoms resilient to ocean alkalinity enhancement: A glimpse of hope?, Sci. Adv., 9, eadg6066, https://doi.org/10.1126/sciadv.adg6066, 2023.

Gore, S., Renforth, P., and Perkins, R.: The potential environmental response to increasing ocean alkalinity for negative emissions, Mitig. Adapt. Strat. Glob. Chang., 24, 1191–1211, https://doi.org/10.1007/s11027-018-9830-z, 2019.

Guo, J. A., Strzepek, R., Willis, A., Ferderer, A., and Bach, L. T.: Investigating the effect of nickel concentration on phytoplankton growth to assess potential side-effects of ocean alkalinity enhancement, Biogeosciences, 19, 3683–3697, https://doi.org/10.5194/bg-19-3683-2022, 2022.

Hartmann, J., West, A. J., Renforth, P., Köhler, P., Rocha, C. L. D. L., Wolf-Gladrow, D. A., Dürr, H. H., and Scheffran, J.: ENHANCED CHEMICAL WEATHERING AS A GEOENGINEERING STRATEGY TO REDUCE ATMOSPHERIC CARBON DIOXIDE, SUPPLY NUTRIENTS, AND MITIGATE OCEAN ACIDIFICATION, Reviews of Geophysics, 1–37, https://doi.org/10.1002/rog.20004, 2013.

Hauck, J., Köhler, P., Wolf-Gladrow, D., and Völker, C.: Iron fertilisation and century-scale effects of open ocean dissolution of olivine in a simulated CO2 removal experiment, Environ. Res. Lett., 11, 024007, https://doi.org/10.1088/1748-9326/11/2/024007, 2016.

Hawco, N. J., McIlvin, M. M., Bundy, R. M., Tagliabue, A., Goepfert, T. J., Moran, D. M., Valentin-Alvarado, L., DiTullio, G. R., and Saito, M. A.: Minimal cobalt metabolism in the marine cyanobacterium *Prochlorococcus*, Proc National Acad Sci, 117, 15740–15747, https://doi.org/10.1073/pnas.2001393117, 2020.

He, J. and Tyka, M. D.: Limits and CO_2_ equilibration of near-coast alkalinity enhancement, Egusphere, 2022, 1–26, https://doi.org/10.5194/egusphere-2022-683, 2022.

Hutchins, D. A. and Boyd, P. W.: Marine phytoplankton and the changing ocean iron cycle, nature climate change, 6, 1072–1079, https://doi.org/10.1038/nclimate3147, 2016.

Hutchins, D. A. and Sañudo-Wilhelmy, S. A.: The Enzymology of Ocean Global Change, Annu Rev Mar Sci, 14, 1–25, https://doi.org/10.1146/annurev-marine-032221-084230, 2021.

Hutchins, D. A., Fu, F.-X., Zhang, Y., Warner, M. E., Feng, Y., Portune, K., Bernhardt, P. W., and Mulholland, M. R.: CO2 control of *Trichodesmium* N_2_ fixation, photosynthesis, growth rates, and elemental ratios: Implications for past, present, and future ocean biogeochemistry, Limnol Oceanogr, 52, 1293–1304, https://doi.org/10.4319/lo.2007.52.4.1293, 2007.

IPCC: IPCC, 2022: Climate Change 2022: Mitigation of Climate Change. Contribution of Working Group III to the Sixth Assessment Report of the Intergovernmental Panel on Climate Change, Popul. Dev. Rev., 48, 629–633, https://doi.org/10.1017/9781009157926, 2022.

Jiang, H.-B., Fu, F.-X., Rivero-Calle, S., Levine, N. M., Sañudo-Wilhelmy, S. A., Qu, P.-P., Wang, X.-W., Pinedo-Gonzalez, P., Zhu, Z., and Hutchins, D. A.: Ocean warming alleviates iron limitation of marine nitrogen fixation, Nat Clim Change, 8, 709–712, https://doi.org/10.1038/s41558-018-0216-8, 2018.

John, S. G., Kelly, R. L., Bian, X., Fu, F., Smith, M. I., Lanning, N. T., Liang, H., Pasquier, B., Seelen, E. A., Holzer, M., Wasylenki, L., Conway, T. M., Fitzsimmons, J. N., Hutchins, D. A., and Yang, S.-C.: The biogeochemical balance of oceanic nickel cycling, Nat Geosci, 15, 906–912, https://doi.org/10.1038/s41561-022-01045-7, 2022.

John, S. G., Mendez, J., Moffett, J., and Adkins, J.: The flux of iron and iron isotopes from San Pedro Basin sediments, Geochim. Cosmochim. Acta, 93, 14–29, https://doi.org/10.1016/j.gca.2012.06.003, 2012.

Karthikeyan, P., Marigoudar, S. R., Nagarjuna, A., and Sharma, K. V.: Toxicity assessment of cobalt and selenium on marine diatoms and copepods, Environ Chem Ecotoxicol, 1, 36–42, doi.org/10.1016/j.enceco.2019.06.001, 2019.

Kazamia, E., Sutak, R., Paz-Yepes, J., Dorrell, R. G., Vieira, F. R. J., Mach, J., Morrissey, J., Leon, S., Lam, F., Pelletier, E., Camadro, J.-M., Bowler, C., and Lesuisse, E.: Endocytosis-mediated siderophore uptake as a strategy for Fe acquisition in diatoms, Sci Adv, 4, eaar4536, https://doi.org/10.1126/sciadv.aar4536, 2018.

Kiran, B., Rani, N., and Kaushik, A.: Environmental toxicity: Exposure and impact of chromium on cyanobacterial species, J Environ Chem Eng, 4, 4137–4142, https://doi.org/10.1016/j.jece.2016.09.021, 2016.

Kling, J. D., Kelly, K. J., Pei, S., Rynearson, T. A., and Hutchins, D. A.: Irradiance modulates thermal niche in a previously undescribed low-light and cold-adapted nano-diatom, Limnol Oceanogr, 66, 2266–2277, https://doi.org/10.1002/lno.11752, 2021.

Lee, M. D., Webb, E. A., Walworth, N. G., Fu, F.-X., Held, N. A., Saito, M. A., and Hutchins, D. A.: Transcriptional Activities of the Microbial Consortium Living with the Marine Nitrogen-Fixing Cyanobacterium *Trichodesmium* Reveal Potential Roles in Community-Level Nitrogen Cycling, Applied and environmental microbiology, 84, e02026-17–16, https://doi.org/10.1128/aem.02026-17, 2018.

Manck, L. E., Park, J., Tully, B. J., Poire, A. M., Bundy, R. M., Dupont, C. L., and Barbeau, K. A.: Petrobactin, a siderophore produced by *Alteromonas*, mediates community iron acquisition in the global ocean, Isme J, 16, 358–369, https://doi.org/10.1038/s41396-021-01065-y, 2022.

Meysman, F. J. R. and Montserrat, F.: Negative CO_2_ emissions via enhanced silicate weathering in coastal environments, Biol Letters, 13, 20160905, https://doi.org/10.1098/rsbl.2016.0905, 2017.

Millero, F. J., Sotolongo, S., and Izaguirre, M.: The oxidation kinetics of Fe(II) in seawater, Geochim Cosmochim Ac, 51, 793–801, https://doi.org/10.1016/0016-7037(87)90093-7, 1987.

Moran, M. A.: The global ocean microbiome, Science, 350, aac8455–aac8455, https://doi.org/10.1126/science.aac8455, 2015.

Morrissey, J. M. and Bowler, C.: Iron utilization in marine cyanobacteria and eukaryotic algae, Frontiers in Microbiology, 3, 1–13, https://doi.org/10.3389/fmicb.2012.00043/abstract, 2012.

Oelkers, E. H., Declercq, J., Saldi, G. D., Gislason, S. R., and Schott, J.: Olivine dissolution rates: A critical review, Chem Geol, 500, 1–19, https://doi.org/10.1016/j.chemgeo.2018.10.008, 2018.

Paasche, E., Brubak, S., Skattebøl, S., Young, J. R., and Green, J. C.: Growth and calcification in the coccolithophorid *Emiliania huxleyi* (Haptophyceae) at low salinities, Phycologia, 35, 394–403, https://doi.org/10.2216/i0031-8884-35-5-394.1, 1996.

Panneerselvam, K., Marigoudar, S. R., and Dhandapani, M.: Toxicity of Nickel on the Selected Species of Marine Diatoms and Copepods, Bull. Environ. Contam. Toxicol., 100, 331–337, https://doi.org/10.1007/s00128-018-2279-7, 2018.

Pettine, M., Millero, F. J., and Noce, T. L.: Chromium (III) interactions in seawater through its oxidation kinetics, Mar Chem, 34, 29–46, https://doi.org/10.1016/0304-4203(91)90012-l, 1991.

Reimers, C. E., Stecher, H. A., Taghon, G. L., Fuller, C. M., Huettel, M., Rusch, A., Ryckelynck, N., and Wild, C.: In situ measurements of advective solute transport in permeable shelf sands, Cont Shelf Res, 24, 183–201, https://doi.org/10.1016/j.csr.2003.10.005, 2004.

Renforth, P. and Henderson, G.: Assessing ocean alkalinity for carbon sequestration, Rev Geophys, 55, 636–674, https://doi.org/10.1002/2016rg000533, 2017.

Rimstidt, J. D., Brantley, S. L., and Olsen, A. A.: Systematic review of forsterite dissolution rate data, Geochim Cosmochim Ac, 99, 159–178, https://doi.org/10.1016/j.gca.2012.09.019, 2012.

Rogelj, J., Popp, A., Calvin, K. V., Luderer, G., Emmerling, J., Gernaat, D., Fujimori, S., Strefler, J., Hasegawa, T., Marangoni, G., Krey, V., Kriegler, E., Riahi, K., Vuuren, D. P. van, Doelman, J., Drouet, L., Edmonds, J., Fricko, O., Harmsen, M., Havlík, P., Humpenöder, F., Stehfest, E., and Tavoni, M.: Scenarios towards limiting global mean temperature increase below 1.5 °C, Nat. Clim. Chang., 8, 325–332, https://doi.org/10.1038/s41558-018-0091-3, 2018.

Rubin, M., Berman-Frank, I., and Shaked, Y.: Dust- and mineral-iron utilization by the marine dinitrogen-fixer *Trichodesmium*, Nature Geoscience, 4, 529–534, https://doi.org/10.1038/ngeo1181, 2011.

Shi, T., Ilikchyan, I., Rabouille, S., and Zehr, J. P.: Genome-wide analysis of diel gene expression in the unicellular N_2_-fixing cyanobacterium *Crocosphaera watsonii* WH 8501, Isme J, 4, 621–632, https://doi.org/10.1038/ismej.2009.148, 2010.

Sunda, W. G. and Huntsman, S. A.: Cobalt and zinc interreplacement in marine phytoplankton: Biological and geochemical implications, Limnol Oceanogr, 40, 1404–1417, https://doi.org/10.4319/lo.1995.40.8.1404, 1995.

Sunda, W. G., Price, N. M., and Morel, F. M. M.: Trace Metal Ion Buffers and Their Use in Culture Studies, Algal Culturing Techniques, 35–63, https://doi.org/10.1016/b978-012088426-1/50005-6, 2005.

Taylor, L. L., Quirk, J., Thorley, R. M. S., Kharecha, P. A., Hansen, J., Ridgwell, A., Lomas, M. R., Banwart, S. A., and Beerling, D. J.: Enhanced weathering strategies for stabilizing climate and averting ocean acidification, Nat. Clim. Chang., 6, 402–406, https://doi.org/10.1038/nclimate2882, 2016.

Tovar-Sanchez, A., Sañudo-Wilhelmy, S. A., Garcia-Vargas, M., Weaver, R. S., Popels, L. C., and Hutchins, D. A.: A trace metal clean reagent to remove surface-bound iron from marine phytoplankton, Mar Chem, 82, 91–99, https://doi.org/10.1016/s0304-4203(03)00054-9, 2003.

Tréguer, P., Bowler, C., Moriceau, B., Dutkiewicz, S., Gehlen, M., Aumont, O., Bittner, L., Dugdale, R., Finkel, Z., Iudicone, D., Jahn, O., Guidi, L., Lasbleiz, M., Leblanc, K., Levy, M., and Pondaven, P.: Influence of diatom diversity on the ocean biological carbon pump, Nat Geosci, 11, 27–37, https://doi.org/10.1038/s41561-017-0028-x, 2018.

Vink, J. P. M. and Knops, P.: Size-Fractionated Weathering of Olivine, Its CO_2_-Sequestration Rate, and Ecotoxicological Risk Assessment of Nickel Release, Mineral-basel, 13, 235, https://doi.org/10.3390/min13020235, 2023.

Weijden, C. H. V. D. and Reith, M.: Chromium(III) — chromium(VI) interconversions in seawater, Mar Chem, 11, 565–572, https://doi.org/10.1016/0304-4203(82)90003-2, 1982.

Welschmeyer, N. A.: Fluorometric analysis of chlorophyll a in the presence of chlorophyll b and pheopigments, Limnol Oceanogr, 39, 1985–1992, https://doi.org/10.4319/lo.1994.39.8.1985, 1994.

Yamamoto, T., Goto, I., Kawaguchi, O., Minagawa, K., Ariyoshi, E., and Matsuda, O.: Phytoremediation of shallow organically enriched marine sediments using benthic microalgae, Mar Pollut Bull, 57, 108–115, https://doi.org/10.1016/j.marpolbul.2007.10.006, 2008.

Yang, N., Lin, Y.-A., Merkel, C. A., DeMers, M. A., Qu, P.-P., Webb, E. A., Fu, F.-X., and Hutchins, D. A.: Molecular mechanisms underlying iron and phosphorus co-limitation responses in the nitrogen-fixing cyanobacterium *Crocosphaera*, Isme J, 16, 2702–2711, https://doi.org/10.1038/s41396-022-01307-7, 2022.

Yoshida, M., Kuma, K., Iwade, S., Isoda, Y., Takata, H., and Yamada, M.: Effect of aging time on the availability of freshly precipitated ferric hydroxide to coastal marine diatoms, Mar. Biol., 149, 379–392, https://doi.org/10.1007/s00227-005-0187-y, 2006.

