## Supplemental Figures for "Responses of globally important phytoplankton species to olivine dissolution products and implications for carbon dioxide removal via ocean alkalinity enhancement"

David A. Hutchins[1†*], Fei-Xue Fu[1], Shun-Chung Yang[1], Seth G. John[1], Stephen J. Romaniello[2], M. Grace Andrews[2], Nathan G. Walworth[1,2,3†*]

[1]University of Southern California, Los Angeles, CA, USA
[2]Vesta, PBC, San Francisco, CA, USA
[3]J. Craig Venter Institute, La Jolla, CA, USA

[†]These authors contributed equally to this work

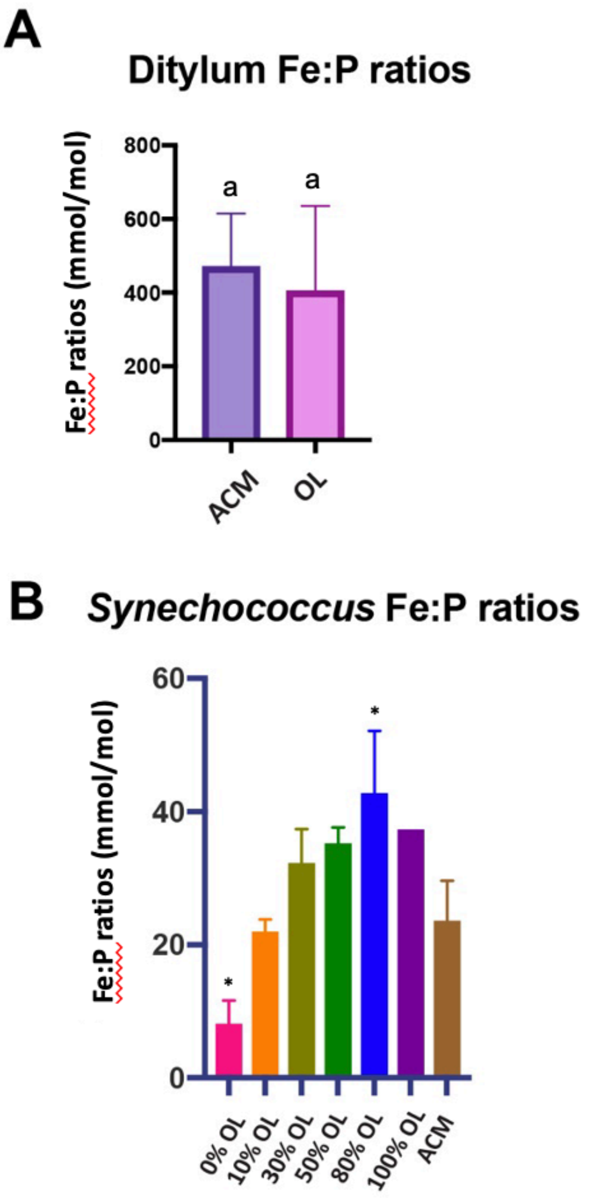


**Supplementary Figure 1. Olivine leachate effects on Fe:P ratios of the diatom *Dytilum* and the picocyanobacterium *Synechococcus.***

**A)** *Dytilum brightwellii* Fe:P ratios; different letters indicate significant differences at the $p < 0.05$ level, and **B)** *Synechococcus* sp. Fe:P ratios in ACM and across a dilution series of OL. Relative to the ACM positive control, significant differences in OL treatments are indicated by $* = p < 0.05$; $** = p < 0.01$; $*** = p < 0.001$; $**** = p < 0.0001$. Abbreviations: OL is olivine leachate, ACM is Aquil control medium. For both species, values represent the means and error bars are the standard deviations of biological triplicate cultures for each treatment.

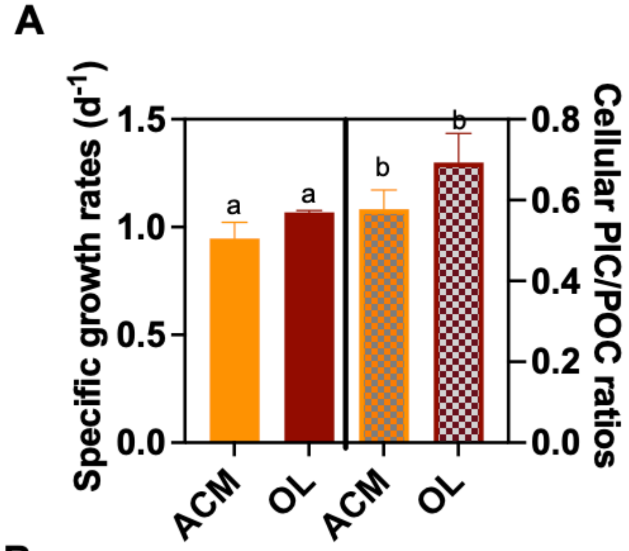


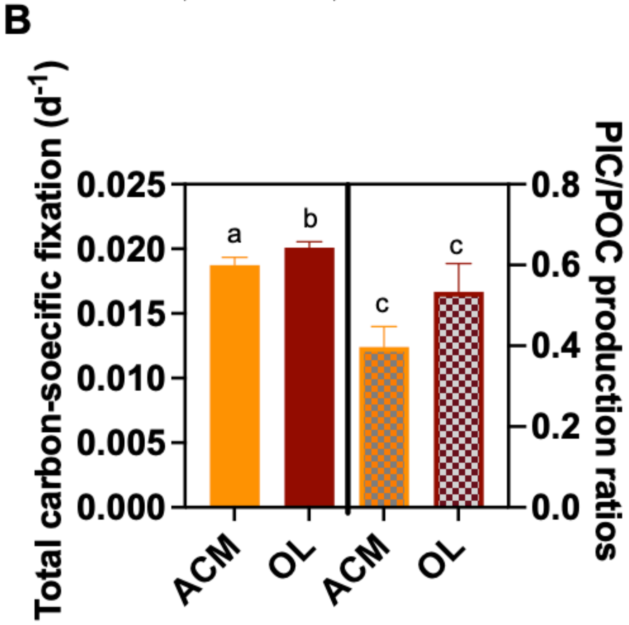


**Supplementary Figure 2. Olivine leachate effects on growth and physiology of a marine coccolithophore.** The coccolithophore *Emiliana huxleyi* was grown in olivine leachate (OL) and in the ACM positive control medium**.** Shown are: **A)** Cell-specific growth rates ($d^{-1}$) on the left (solid bars) and cellular PIC/POC ratios (mol:mol, calcite/organic carbon) on the right (hatched bars). **B)** Carbon-specific fixation rates ($hr^{-1}$) on the left (solid bars) and PIC:POC production ratios (calcification rate/organic carbon production rate) on the right (hatched bars). Values represent the means and error bars are the standard deviations of biological triplicate cultures for each treatment. Different letters indicate significant differences at the $p < 0.05$ level.

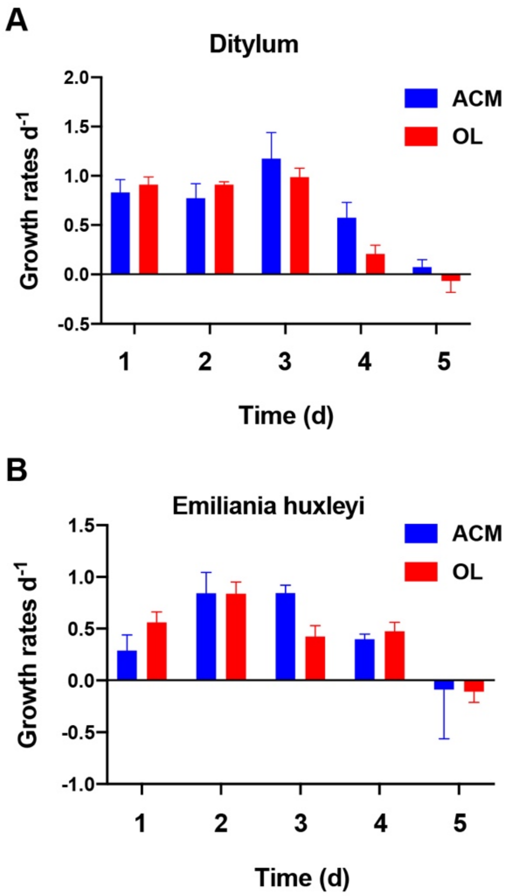


**Supplementary Figure 3. Effects of olivine leachate versus culture medium controls on growth competition during co-culture of a diatom and a coccolithophore.** Shown are trends in specific growth rates ($d^{-1}$) over 5 days of co-culture for **A)** the diatom *Ditylum brightwellii* and **B)** the coccolithophore *Emiliana huxleyi* grown in olivine leachate (OL, red symbols) and ACM positive control medium (blue symbols). Values represent the means and error bars are the standard deviations of biological triplicate cultures for each treatment.
